# Introducing RpsA Point Mutation Δ438A or D123A into the Chromosome of *M. tuberculosis* Confirms its Role in Causing Resistance to Pyrazinamide

**DOI:** 10.1101/505297

**Authors:** Wanliang Shi, Peng Cui, Hongxia Niu, Shuo Zhang, Tone Tønjum, Bingdong Zhu, Ying Zhang

## Abstract

Pyrazinamide (PZA) is a unique frontiline drug for shortening tuberculosis treatment, but its mechanisms of action are elusive. We previously identified RpsA as a target of PZA and found an alanine deletion at position 438 (Δ438A) in RpsA associated with PZA resistance, but its role in PZA resistance is controversial. Here, we introduced RpsA mutation Δ438A or D123A into *M. tuberculosis* chromosome and demonstrated that the introduced RspA mutations are indeed responsible for PZA resistance.

## TEXT

PZA is an important first-line drug that plays a unique role in shortening TB therapy from 9-12 months to 6 months due to its unique sterilizing activity in killing persisters that are not killed by other TB drugs (7, 29). Despite its high activity in vivo, PZA is a peculiar drug that has virtually no activity in vitro under normal culture conditions (19, 29), but is active at acid pH (6, 31). Recent studies of new drugs such as bedaquiline, PA-824, and moxifloxacin have highlighted the essentiality of PZA as these new agents mainly work in conjunction with PZA but cannot replace it (20, 21). Thus, PZA is a key drug that will likely play a major role in new drug regimens for treatment of TB and drug-resistant TB.

It is well established that mutations in the *pncA* gene encoding nicotinamidase/pyrazinamidase (PZase) involved in conversion of PZA to active form pyrazinoic acid (POA) are the major mechanism of PZA resistance (11, 32), accounting for85% of all PZA resistance (32). However, some PZA-resistant strains without *pncA* mutations may have mutations in potential targets of PZA, including ribosomal protein S1 (RpsA) involved in trans-translation (5, 9, 13, 17, 24), aspartate decarboxylase (PanD) (12, 27) involved in synthesis of β-alanine – a precursor for pantothenate and CoA biosynthesis, and ATP-dependent protease ClpC1(26, 28). We have previously identified a 3–base pair “GCC” deletion resulting in loss of an Alanine at amino acid 438 in RpsA in a low-level PZA-resistant clinical isolate DHM444 (MIC = 200 to 300 μg/ml PZA compared with 100 μg/ml in susceptible *M. tuberculosis)* without *pncA* mutation (13). *rpsA* mutations are subsequently shown to be associated with PZA resistance in clinical strains (3, 14, 18). However, certain nonsynonymous mutations in RpsA protein (A364G) seem to occur in some PZA susceptible strains (1). In addition, a recent study by Dillon et al. claims that RpsA is not a target of PZA and that the RpsA Δ438A mutation does not cause PZA resistance (2). These discordant results raise doubts about RpsA being a target of PZA and the role of RpsA Δ438A mutation in PZA resistance.

In this study, to more convincingly address the role of the RpsA Δ438A mutation in PZA resistance, we transferred this mutation into the genome of *M. tuberculosis* H37Rv by homologous recombination. It is worth noting that the same strategy has been successfully used to demonstrate *inhA* and certain *embB* point mutations being responsible for INH and ethambutol resistance, respectively (10, 22). The RpsA Δ438A mutation was created by a two-step allelic exchange method as described (8). Briefly, a 3,440-bp fragment spanning the *rpsA* Δ438A deletion mutation was amplified by PCR with DHM444 genomic DNA using primers containing HindIII and PacI restriction sites (underlined) in the forward primer (FrpsA 5’-CGG<u>AAGCTT</u>CCACACCACGTTCAACCAGAC-3’ and reverse primer RrpsA 5’-GC<u>TTAA</u>TTAAGCACGCGCTTGTGCCACAGAG-3’), respectively. The PCR fragment was then cloned into the p2NIL vector followed by insertion of a PacI cassette containing the *sacB* and *lacZ.* The recombinant plasmid was transformed in *M. tuberculosis* H37Rv as described (12). The desired sucrose-resistant but kanamycin-susceptible transformants were analyzed by PCR and sequenced to confirm that the transformed *M. tuberculosis* has the correct RpsA Δ438A mutation (Fig. 1). We determined the PZA MICs for the RpsA Δ438A mutant and the parent strain *M. tuberculosis* H37Rv using the proportion method and found them to be 200 μg/ml and 100 μg/ml (pH5.8), respectively. In addition, we measured the PZA and POA MICs for the RpsA Δ438A mutant in liquid 7H9 medium (pH5.8) using microdilution method. The PZA MIC for the *M. tuberculosis* RpsA Δ438A mutant strain was 300 μg/ml while the parent strain H37Rv was susceptible at this concentration (Fig. 2A). The POA MIC for the *M. tuberculosis* RpsA Δ438A mutant strain was 50 μg/ml compared with 25 μg/ml for the parent strain H37Rv (Fig. 2B). On the other hand, both strains had the same MIC for the control drugs isoniazid (INH) (0.03 μg/ml) and rifampin (RIF) (0.12 μg/ml) (Fig. 2C, 2D). These results clearly demonstrate that transfer of the RpsA Δ438A mutation into the genome of *M. tuberculosis* caused resistance to PZA and POA specifically, but not to INH and RIF. This provides conclusive evidence that the RpsA Δ438A mutation is indeed responsible for the PZA resistance in the strain DHM444.

**Fig. 1.**
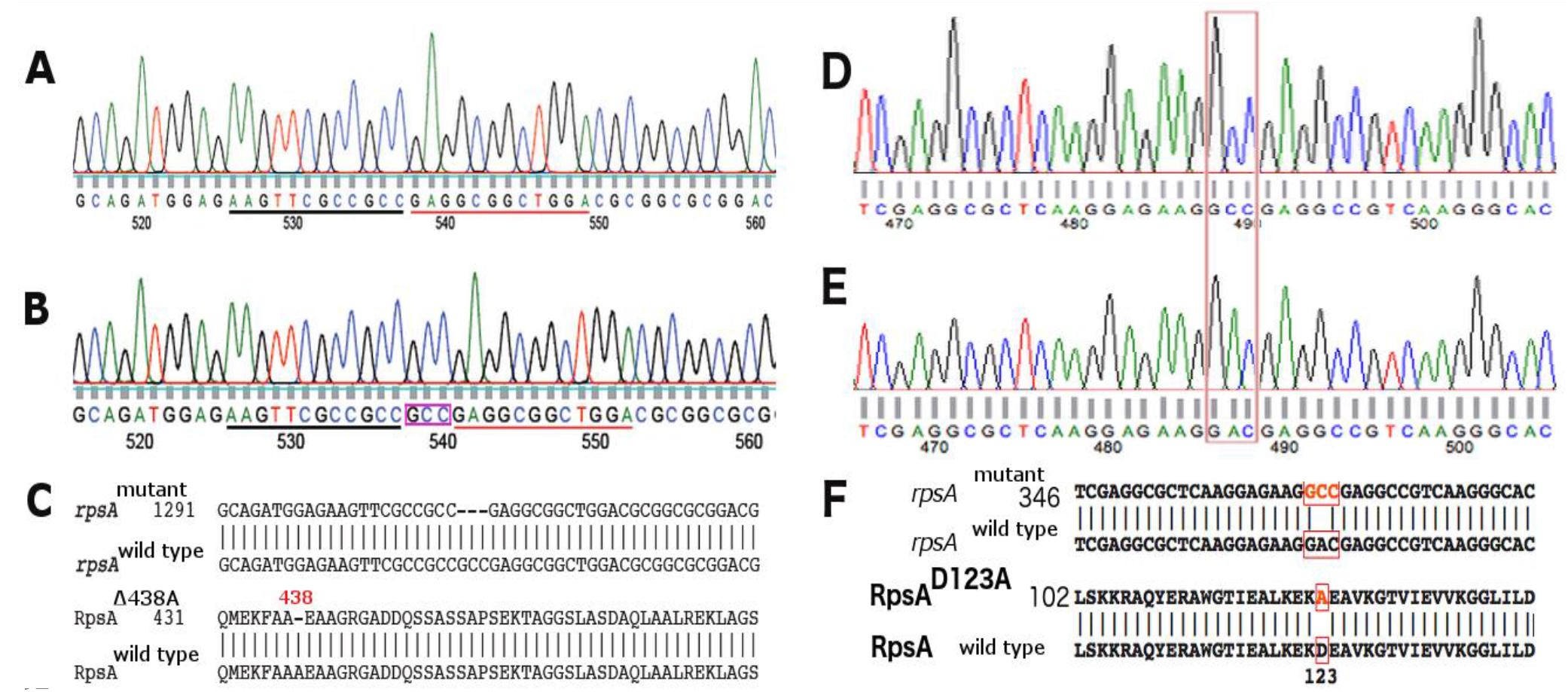
Confirmation of RpsA Δ438A and D123A point mutation construction by Sanger sequencing. **A**. Chromatogram of partial *rpsA* sequence showing the “GCC” (coding for Alanine, A) deletion in the constructed RpsAΔ438A mutant from PZA-resistant strain DHM444. **B**. Chromatogram of partial *rpsA* sequence showing the presence of “GCC” (Alanine) in the *M. tuberculosis* H37Rv parent strain. **C**. Alignments of partial *rpsA* sequence from *M. tuberculosis* H37Rv wild type and the constructed RpsA Δ438A mutant at the nucleotide level (upper panel) and amino acid level (lower panel). D and E. Chromatogram of partial *rpsA* sequence showing “GAC” (wild type) change to “GCC” which is contained in one PZA-resistant clinical isolate. **F**. Alignments of partial *rpsA* sequence from *M. tuberculosis* H37Rv wild type and the constructed RpsA D123A mutant at the nucleotide level (upper panel) and amino acid level (lower panel).

**Fig. 2.**
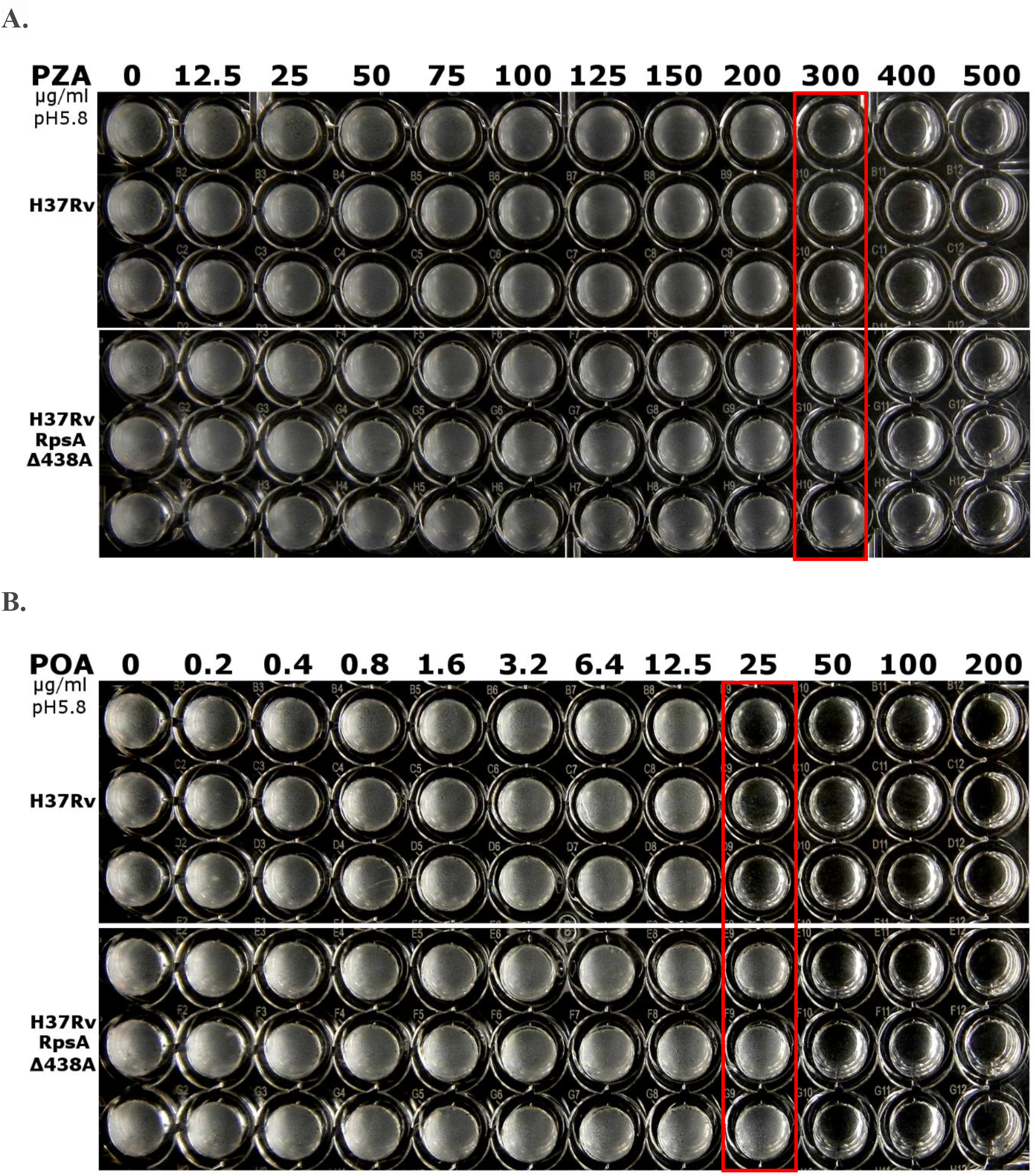
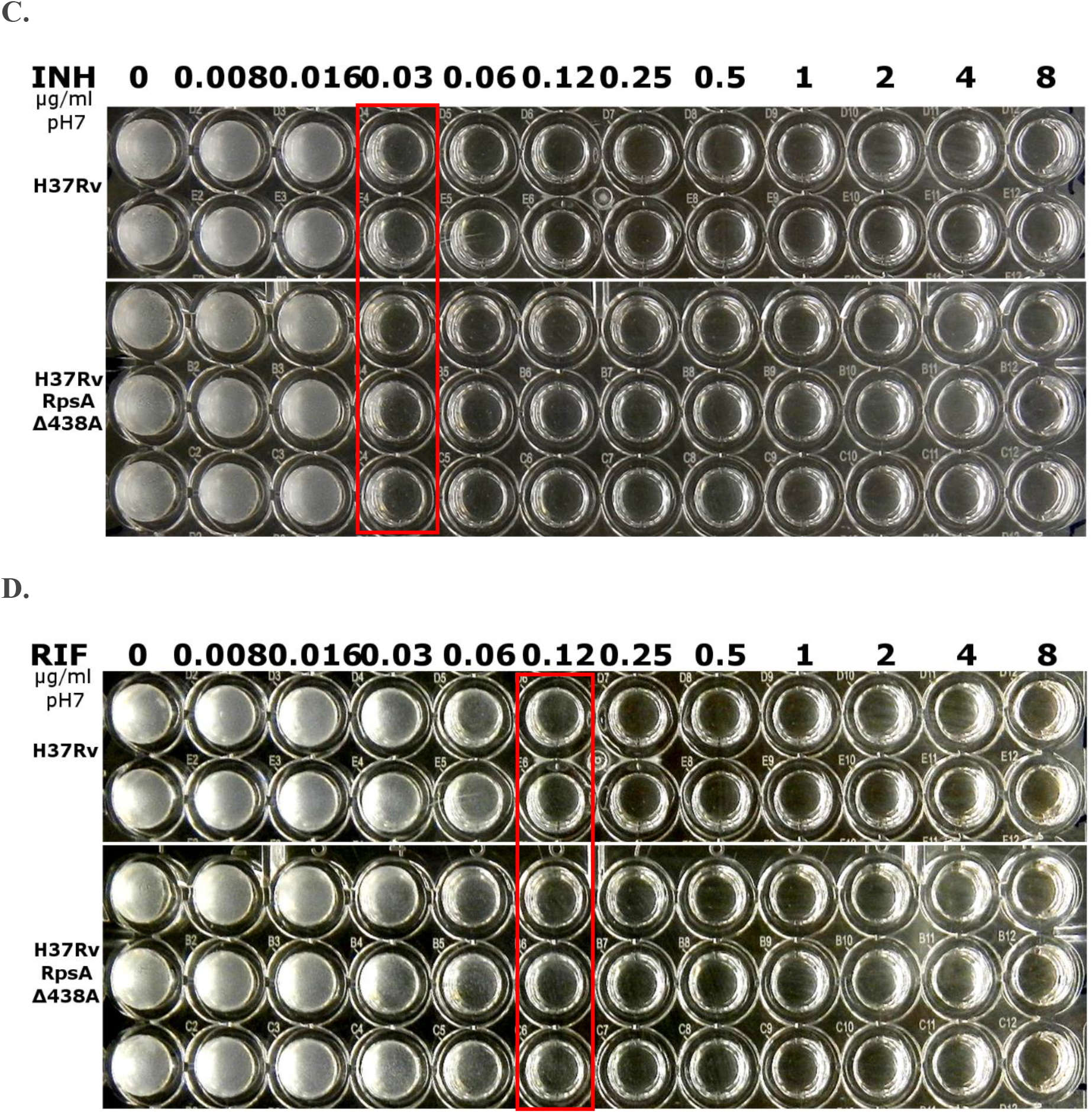
Drug susceptibility testing (DST) results of the constructed RpsA Δ438A mutant strain compared with the control strain H37Rv. The drug susceptibility testing was performed in 7H9/ADC broth using microdilution method with pH5.8 for PZA and POA DST, and pH 7.0 for INH and RIF DST. The constructed RpsA Δ438A mutant strain was more resistant to PZA (A) and POA (B) than the control strain H37Rv but was as susceptible to INH (C) and RIF (D) as the control strain H37Rv. **A**. The PZA MIC of the *M. tuberculosis* H37Rv control strain was 300 μg/ml while the *M. tuberculosis* RpsA Δ438A mutant strain was resistant at this concentration. **B**. The POA MIC of *M. tuberculosis* H37Rv parent strain was 25 μg/ml while the POA MIC for the *M. tuberculosis* H37Rv RpsA Δ438A mutant strain was 50 μg/ml. **C**. Isoniazid (INH) MICs for the parent strain *M. tuberculosis* H37Rv and the *M. tuberculosis* H37Rv RpsA Δ438A mutant were the same (0.03 μg/ml). **D**. The RIF MICs for the parent strain *M. tuberculosis* H37Rv and the *M. tuberculosis* H37Rv RpsA Δ438A mutant were the same (0.12 μg/ml). The red boxes are used to highlight the difference in PZA and POA susceptibility between the RpsA Δ438A mutant strain and the parent strain H37Rv.

In addition, in our previous study we found another low level PZA-resistant clinical isolate that has two *rpsA* mutations RpsA T5S and D123A with wild type *pncA* gene (13). To provide further proof that RpsA point mutations can cause PZA resistance, we constructed another RpsA mutation D123A into the genome of *M. tuberculosis* H37Rv. Site-directed mutagenesis of *rpsA* was performed with primers S1D123AF 5’-GCTCAAGGAGAAGGCCGAGGCCGTCAAGG-3’and S1D123AR 5’-CCTTGACGGCCTCGGCCTTCTCCTTGAGC-3’using the QuikChange™ site-directed mutagenesis kit as described by the manufacturer (Agilent Technologies). Then the point mutation RpsA D123A was transferred to the parental strain *M. tuberculosis* H37Rv as decribed as above. We found that indeed the PZA MIC for the *M. tuberculosis* RpsA D123A mutant was increased by 2 fold (200 μg/ml PZA) compared with the parental strain *M. tuberculosis* H37Rv (100 μg/ml PZA) on 7H11 agar plate at pH5.8.

It is worth discussing the possible causes for the discrepant results between the Dillon et al. study (2) and ours. One possibility is the unusual and not accepted method used by that group to determine the PZA susceptibility by using OD600 and MIC90. PZA susceptibility testing is notoriously difficult because of the acid pH requirement for drug activity and many factors affect the susceptibility results (30). The inability to observe PZA or POA resistance in their *rpsA* overexpression or point mutation strains could be attributed to the questionable method used in that study. Another argument used by Dillon et al. against the RpsA Δ438A mutation conferring PZA resistance (2) is that DHM444’s PZA resistance could be due to defective PZase activity reported in a previous study (16). To address this question, we tested PZase activity for DHM444 as described (27) and found that contrary to the previous report (2), The PZA-resistant strain DHM444 still has PZase activity (Data not shown). Thus, the low level PZA resistance in DHM444 could not be attributed to defective PZase activity, and this study provides clear evidence that its PZA resistance is most likely due to the RpsA Δ438A mutation (Fig. 2A, 2B).

That RpsA is a target of PZA (13) is supported by a number of observations including association of RpsA mutations and PZA resistance in clinical strains (3, 5, 9, 14, 18, 23), *rpsA* overexpression causing PZA resistance (13), as well as biochemical and structural studies showing the binding of the drug to the RpsA (4, 15, 25). Despite these supporting studies, and in view of the question about the role of RpsAΔ438A mutation in PZA resistance, point mutation construction into the chromosome, though challenging, is by far the most convincing method to prove if a specific mutation is the cause for drug resistance. Our findings that the RpsAΔ438A mutation and the D123A mutation when introduced into the *M. tuberculosis* chromosome are indeed responsible for PZA resistance provide further proof that RpsA is a real target of PZA. Future studies are needed to address the role of certain *rpsA* mutations in some seemingly “PZA-susceptible” strains by constructing more isogenic strains with *rpsA* mutations to determine whether they are involved in PZA resistance while others may not or may confer only borderline resistance that is easily mischaracterized as PZA susceptible due to insensitive PZA DST.

## ACKNOWLEDGMENTS

This work was supported by NIH R01AI099512 and AMR-PART, Research Council of Norway.

## REFERENCES

1. Alexander, D. C., J. H. Ma, J. L. Guthrie, J. Blair, P. Chedore, and F. B. Jamieson. 2012. Gene sequencing for routine verification of pyrazinamide resistance in Mycobacterium tuberculosis: a role for pncA but not rpsA. Journal of clinical microbiology 50:3726–8.

2. Dillon, N. A., N. D. Peterson, H. A. Feaga, K. C. Keiler, and A. D. Baughn. 2017. Anti-tubercular Activity of Pyrazinamide is Independent of trans-Translation and RpsA. Sci Rep 7:6135.

3. Gu, Y., X. Yu, G. Jiang, X. Wang, Y. Ma, Y. Li, and H. Huang. 2016. Pyrazinamide resistance among multidrug-resistant tuberculosis clinical isolates in a national referral center of China and its correlations with pncA, rpsA, and panD gene mutations. Diagnostic microbiology and infectious disease 84:207–11.

4. Huang, B., J. Fu, C. Guo, X. Wu, D. Lin, and X. Liao. 2016. (1)H, (15)N, (13)C resonance assignments for pyrazinoic acid binding domain of ribosomal protein S1 from Mycobacterium tuberculosis. Biomol NMR Assign 10:321–4.

5. Khan, M. T., S. I. Malik, A. I. Bhatti, S. Ali, A. S. Khan, M. T. Zeb, T. Nadeem, and S. Fazal. 2018. Pyrazinamide-resistant mycobacterium tuberculosis isolates from Khyber Pakhtunkhwa and rpsA mutations. J Biol Regul Homeost Agents 32:705–709.

6. McDermott, W., and R. Tompsett. 1954. Activation of pyrazinamide and nicotinamide in acidic environment in vitro. Am. Rev. Tuberc. 70:748–754.

7. Mitchison, D. A. 1985. The action of antituberculosis drugs in short-course chemotherapy. Tubercle 66:219–25.

8. Parish, T., and N. G. Stoker. 2000. Use of a flexible cassette method to generate a double unmarked Mycobacterium tuberculosis tlyA plcABC mutant by gene replacement. Microbiology 146 (Pt 8):1969–75.

9. Ramirez-Busby, S. M., T. C. Rodwell, L. Fink, D. Catanzaro, R. L. Jackson, M. Pettigrove, A. Catanzaro, and F. Valafar. 2017. A Multinational Analysis of Mutations and Heterogeneity in PZase, RpsA, and PanD Associated with Pyrazinamide Resistance in M/XDR Mycobacterium tuberculosis. Sci Rep 7:3790.

10. Safi, H., B. Sayers, M. H. Hazbon, and D. Alland. 2008. Transfer of embB codon 306 mutations into clinical Mycobacterium tuberculosis strains alters susceptibility to ethambutol, isoniazid, and rifampin. Antimicrobial Agents and Chemotherapy 52:2027–34.

11. Scorpio, A., and Y. Zhang. 1996. Mutations in pncA, a gene encoding pyrazinamidase/nicotinamidase, cause resistance to the antituberculous drug pyrazinamide in tubercle bacillus. Nat Med 2:662–7.

12. Shi, W., J. Chen, J. Feng, P. Cui, S. Zhang, X. Weng, W. Zhang, and Y. Zhang. 2014. Aspartate decarboxylase (PanD) as a new target of pyrazinamide in Mycobacterium tuberculosis. Emerg Microbes Infect 3:e58.

13. Shi, W., X. Zhang, X. Jiang, H. Yuan, J. S. Lee, C. E. Barry, 3rd, H. Wang, W. Zhang, and Y. Zhang. 2011. Pyrazinamide inhibits trans-translation in *Mycobacterium tuberculosis*. Science 333:1630–2.

14. Simons, S. O., A. Mulder, J. van Ingen, M. J. Boeree, and D. van Soolingen. 2013. Role of rpsA Gene Sequencing in Diagnosis of Pyrazinamide Resistance. Journal of clinical microbiology 51:382.

15. Singh, A., P. Somvanshi, and A. Grover. 2018. Pyrazinamide drug resistance in RpsA mutant (438A) of Mycobacterium tuberculosis: Dynamics of essential motions and free-energy landscape analysis. J Cell Biochem.

16. Speirs, R. J., J. T. Welch, and M. H. Cynamon. 1995. Activity of n-propyl pyrazinoate against pyrazinamide-resistant Mycobacterium tuberculosis: investigations into mechanism of action of and mechanism of resistance to pyrazinamide. Antimicrob Agents Chemother 39:1269–71.

17. Tan, Y., Z. Hu, T. Zhang, X. Cai, H. Kuang, Y. Liu, J. Chen, F. Yang, K. Zhang, S. Tan, and Y. Zhao. 2014. Role of pncA and rpsA gene sequencing in detection of pyrazinamide resistance in Mycobacterium tuberculosis isolates from southern China. J Clin Microbiol 52:291–7.

18. Tan, Y., Z. Hu, T. Zhang, X. Cai, H. Kuang, Y. Liu, J. Chen, F. Yang, K. Zhang, S. Tan, and Y. Zhao. 2014. Role of pncA and rpsA gene sequencing in detection of pyrazinamide resistance in Mycobacterium tuberculosis isolates from southern China. Journal of clinical microbiology 52:291–7.

19. Tarshis, M. S., and W. A. Weed, Jr. 1953. Lack of significant in vitro sensitivity of Mycobacterium tuberculosis to pyrazinamide on three different solid media. Am. Rev. Tuberc. 67:391–5.

20. Tasneen, R., S. Y. Li, C. A. Peloquin, D. Taylor, K. N. Williams, K. Andries, K. E. Mdluli, and E. L. Nuermberger. 2011. Sterilizing Activity of Novel TMC207-and PA-824-Containing Regimens in a Murine Model of Tuberculosis. Antimicrobial Agents and Chemotherapy 55:5485–92.

21. Tasneen, R., S. Tyagi, K. Williams, J. Grosset, and E. Nuermberger. 2008. Enhanced bactericidal activity of rifampin and/or pyrazinamide when combined with PA-824 in a murine model of tuberculosis. Antimicrob Agents Chemother 52:3664–8.

22. Vilcheze, C., F. Wang, M. Arai, M. H. Hazbon, R. Colangeli, L. Kremer, T. R. Weisbrod, D. Alland, J. C. Sacchettini, and W. R. Jacobs, Jr. 2006. Transfer of a point mutation in Mycobacterium tuberculosis inhA resolves the target of isoniazid. Nat Med 12:1027–9.

23. Werngren, J., E. Alm, and M. Mansjo. 2017. Non-pncA Gene-Mutated but Pyrazinamide-Resistant Mycobacterium tuberculosis: Why Is That? Journal of clinical microbiology 55:1920–1927.

24. Werngren, J., E. Alm, and M. Mansjo. 2017. Non pncA gene-mutated but pyrazinamide resistant M. tuberculosis, why is that? J Clin Microbiol.

25. Yang, J., Y. Liu, J. Bi, Q. Cai, X. Liao, W. Li, C. Guo, Q. Zhang, T. Lin, Y. Zhao, H. Wang, J. Liu, X. Zhang, and D. Lin. 2015. Structural basis for targeting the ribosomal protein S1 of Mycobacterium tuberculosis by pyrazinamide. Molecular microbiology 95:791–803.

26. Yee, M., P. Gopal, and T. Dick. 2017. Missense Mutations in the Unfoldase ClpC1 of the Caseinolytic Protease Complex Are Associated with Pyrazinamide Resistance in Mycobacterium tuberculosis. Antimicrob Agents Chemother 61. DOI: 10.1128/AAC.02342-16

27. Zhang S, Chen J, Shi W, Liu W, Zhang WH, and Z. Y. 2013. Mutations in panD encoding aspartate decarboxylase are associated with pyrazinamide resistance in Mycobacterium tuberculosis. Emerging Microbes & Infections 2:e34; doi:10.1038/emi.2013.38.

28. Zhang, S., J. Chen, W. Shi, P. Cui, J. Zhang, S. Cho, W. Zhang, and Y. Zhang. 2017. Mutation in clpC1 encoding an ATP-dependent ATPase involved in protein degradation is associated with pyrazinamide resistance in Mycobacterium tuberculosis. Emerg Microbes Infect 6:e8.

29. Zhang, Y., and D. Mitchison. 2003. The curious characteristics of pyrazinamide: a review. Int J Tuberc Lung Dis 7:6–21.

30. Zhang, Y., S. Permar, and Z. Sun. 2002. Conditions that may affect the results of susceptibility testing of Mycobacterium tuberculosis to pyrazinamide. J Med Microbiol 51:42–49.

31. Zhang, Y., A. Scorpio, H. Nikaido, and Z. Sun. 1999. Role of acid pH and deficient efflux of pyrazinoic acid in unique susceptibility of Mycobacterium tuberculosis to pyrazinamide. J Bacteriol 181:2044–2049.

32. Zhang, Y., W. Shi, W. Zhang, and D. Mitchison. 2014. Mechanisms of Pyrazinamide Action and Resistance. Microbiol Spectrum, ASM Press 2(4):July 2014, 2:2.4.03. doi:10.1128/microbiolspec.MGM2-0023-2013.

